# Metastatic breast cancer cells induce altered microglial morphology and electrical excitability *in vivo*

**DOI:** 10.1101/636159

**Authors:** Anna Simon, Ming Yang, Joanne L. Marrison, Andrew D. James, Peter J. O’Toole, Paul M. Kaye, Miles A. Whittington, Sangeeta Chawla, William J. Brackenbury

**Author notes:** Corresponding authors: Dr Sangeeta Chawla and Dr William J. Brackenbury, Department of Biology and York Biomedical Research Institute, University of York, Wentworth Way, Heslington, York YO10 5DD, UK.

## Abstract

**Background:** An emerging problem in the treatment of breast cancer is the increasing incidence of metastases to the brain. Metastatic brain tumours are incurable and can cause epileptic seizures and cognitive impairment, so better understanding of this niche, and the cellular mechanisms, is urgently required. Microglia are the resident brain macrophage population, becoming “activated” by neuronal injury, eliciting an inflammatory response. Microglia promote proliferation, angiogenesis and invasion in brain tumours and metastases. However, the mechanisms underlying microglial involvement appear complex and better models are required to improve understanding of function.

**Methods:** Here, we sought to address this need by developing a model to study metastatic breast cancer cell-microglial interactions using intravital imaging combined with *ex vivo* electrophysiology. We implanted an optical window on the parietal bone to facilitate observation of cellular behaviour *in situ* in the outer cortex of heterozygous *Cx3cr1*^GFP/+^ mice. Results: We detected GFP-expressing microglia in *Cx3cr1*^GFP/+^ mice up to 350 µm below the window without significant loss of resolution. When DsRed-expressing metastatic MDA-MB-231 breast cancer cells were implanted in Matrigel under the optical window, significant accumulation of activated microglia around invading tumour cells could be observed. This inflammatory response resulted in significant cortical disorganisation and aberrant spontaneously-occurring local field potential spike events around the metastatic site.

**Conclusions:** These data suggest that peritumoral microglial activation and accumulation may play a critical role in local tissue changes underpinning aberrant cortical activity, which offers a possible mechanism for the disrupted cognitive performance and seizures seen in patients with metastatic breast cancer.

## Introduction

Metastasis, the spreading of primary tumours to secondary sites, is responsible for 90 % of deaths from cancer and is a leading cause of morbidity in cancer patients (1). The incidence of breast cancer patients developing brain metastases is increasing, particularly for those with tumours that are human epidermal growth factor 2 (HER2) positive or triple negative (TN; lacking HER2, progesterone and oestrogen receptor) (2, 3). Most breast cancer brain metastases occur in the frontal lobe, with cerebellum and parietal lobe also being common sites (4). The majority are characterised as parenchymal brain metastasis, with 10-20 % occurring as leptomeningeal metastasis (5). Treatment options are mainly limited to whole brain radiation, in some cases neurosurgery, and survival is poor (6, 7). Furthermore, patients with metastatic brain tumours present with neurological symptoms including cognitive impairment and epileptic seizures (8, 9), and this can have a significant impact on quality of life. However, the pathophysiological mechanisms underlying the cause of neurocognitive impairment and tumour-associated epilepsy are poorly understood (10-12). Thus, there is an urgent need to better understand the mechanisms regulating seeding and expansion of brain metastases and their neurological sequelae in order to develop new effective therapies.

As is the case with primary tumours, the heterogeneous microenvironment is thought to play a key role in progression of metastatic disease (13). In the context of the neural niche, the heterotypic contributions of a range of cell types has been identified, including microglia. Microglia are the resident innate immune (macrophage) population in the brain and play a key role in neuroinflammatory processes. At steady state, microglia undertake a surveillance role, becoming “activated” following neuronal injury or infection (14). Microglial activation drives the neuroinflammatory responses that underpin neuronal loss and dysfunction in neurodegeneration, affective disorders, stroke and brain tumours (15-18). Under normal physiological conditions *in vivo*, microglia adopt a highly mobile ramified morphology, constantly scanning the brain parenchyma for injury to neuronal circuitry. Upon detection of a lesion, activated microglia rapidly undergo targeted migration to the injury site where they orchestrate an inflammatory response (19). Activated microglia are the main immune population clustering around brain metastases in mouse models and patient brain tissue (20).

The involvement of microglia in progression of metastases is complex and dependent on programming by the local brain tumour microenvironment (18). Microglia interact with metastatic breast cancer cells in a heterogeneous fashion in mouse models, adopting multiple morphological states related to diverse functional contributions to tumour progression (20). The spectrum of activated microglial phenotypes in brain tumours ranges from “classical” M1 proinflammatory cells capable of releasing proinflammatory cytokines and eliciting a T cell response, to “alternative” M2 anti-inflammatory cells which promote tumour growth, angiogenesis and invasion (21-25). Thus, targeting the anti-inflammatory microglial population (26-28), and/or enhancing the activity of the proinflammatory population (29), may provide a new treatment approach for brain metastases. However, better models are required to improve understanding of the functional activity of microglia in the microenvironment around brain tumour cells.

Intravital two-photon imaging represents a useful tool for studying cellular activity in the microenvironment of tumours in three dimensions and over time (30). Using this technique, it is possible to visualise tissue structures at cellular resolution in order to understand mechanisms underlying cell-cell interactions and invasion. Intravital imaging approaches using orthotopic breast tumour models have provided important insights into the mechanisms underlying migration and invasion as well as improving understanding of tumour heterogeneity (31-34). In the context of brain tumours, intravital imaging has been used to visualise glioma proliferation and invasion, as well as response to anti-tumour therapies (35). Intravital imaging has also been employed to study microglial activation (14, 36). However, the approach has not been previously used to directly study the interaction between breast cancer cells and microglia in the brain metastatic niche. In the present study, we sought to address this gap by developing a model to study breast tumour-microglial interactions using intravital imaging combined with *ex vivo* electrophysiology. Using this approach, it was possible to visualise microglial activation and accumulation, cortical disorganisation and aberrant spontaneous electrical activity around the lesion site.

## Materials and Methods

### Cell culture

MDA-MB-231 cells were maintained in Dulbecco’s modified eagle medium (DMEM) supplemented with 5 % FBS and 4 mM L-glutamine (37). The cells were stably transduced with recombinant lentivirus for DsRed (gift from M. Lorger, University of Leeds). Cells were confirmed as mycoplasma-free using the DAPI method and their molecular identity was confirmed by short tandem repeat analysis.

### Animals

Female heterozygous *Cx3cr1*^GFP/+^ mice on a C57BL/6 (B6) background (obtained from the Jackson Laboratories, Bar Harbour, USA) were used for all experiments in this study (38). Animals were maintained on a 12 h light/dark cycle with food/water *ad libitum*. Animal procedures were performed following approval from the University of York Animal Welfare and Ethical Review Body and under authority of a UK Home Office Project Licence. Surgical procedures were performed within the regulations of the UK Animals (Scientific Procedures) Act, 1986. Experiments have been reported in accordance with the ARRIVE guidelines.

### Cranial window installation and implantation of tumour cells

The installation of a cranial imaging window was based on procedures described previously (39). Mice (8-17 weeks old; n = 23) were anaesthetised using isoflurane and the head was stabilised in a stereotaxic frame (Kopf Instruments). Cerebral oedema was treated peri-operatively with dexamethasone (2 µg/g body weight; intramuscular injection). Post-surgical inflammation was treated peri-operatively and at 24 h intervals for 3 days followed by 48 h intervals for 2 weeks after surgery with carprofen (5 µg/g body weight; subcutaneous injection) and buprenorphine (0.1 µg/g body weight; subcutaneous injection). A small (3 mm diameter) craniotomy was performed on the left parietal bone above the posterior parietal cortex (centred 2.5 mm posterior to Bregma, 2 mm lateral to midline). In some animals, MDA-MB-231-DsRed cells (10^4^ suspended in 20 % v/v Matrigel in saline) were placed in a single drop on the exposed area using a pipette. A 4 mm diameter round No. 1 glass coverslip (Menzel-Gläser) was then placed onto the surgical area, mounted onto the surrounding skull using Vetbond tissue adhesive and sealed in place with dental cement. A small 4 mm × 8 mm titanium plate was attached to the skull using tissue adhesive as an aid for future stabilisation of the head during imaging. Animals were allowed to recover for 5 days before imaging. Weight and body condition were monitored at regular intervals following surgery. Mice were re-grouped following recovery from surgery and were maintained for varying durations up to 1 month following surgery to allow for serial imaging.

### Microscope set-up

We used a Zeiss LSM 780 multiphoton fitted to an AxioObserver Z1 inverted microscope and coupled to a Coherent Chameleon Ultra Ti-Sapphire 2-photon laser for this study with the stage sitting within a Solent Scientific temperature-controlled light-tight box. An InverterScope (LSM Tech, Etters, PA) objective inverter was used to convert the system to an upright configuration and enable positioning of the W Plan Apochromat 20x/1.0 NA objective lens above the cranial window. The inverter was purpose adapted, with the addition of a reflective interior and MaxMirror dielectric mirrors, to maximise excitation and emission transmission at both visible and far-red wavelengths (>900 nm). The excitation power exiting a Plan Apochromat 20×/0.8 objective was monitored *a priori* using a Coherent Fieldmate power meter and Coherent OP-2 vis sensor (488 nm) or a PM2 sensor (920 nm) for a range of % transmissions with and without the InverterScope. A standard control sample (Molecular Probes FluorCell slide #6) was used to compare Alexafluor 488 emission back through to the detectors with and without the InverterScope after excitation at either 488 nm or 920 nm.

### Intravital imaging

Mice were anaesthetised for restraint using ketamine (100 µg/g body weight) and xylazine (10 µg/g body weight) prior to imaging and placed on their ventral side on the heated microscope stage. The head was stabilised with an in-house custom-made holder to minimise motion artefacts. The light-tight box was pre-warmed to 37 °C and was maintained at this temperature for the duration of the imaging session (typically less than 1 h). GFP and DsRed were visualised using two-photon excitation at 920 nm and emission detected via a 500-550 nm band pass filter for GFP and 550 nm long pass filter for DsRed. Images were acquired sequentially and saved for offline processing using Zeiss Zen2 software. The laser power used was up to 15 mW post-objective, dependent on experiment.

### Electrophysiology

Coronal slices (450 µm thick) were prepared from the region of the brain under the imaging window, and the contralateral equivalent, at the end of the intravital imaging period. Slices were maintained in an interface chamber at 34 °C in oxygenated (95 % O_2_/5 % CO_2_) artificial cerebrospinal fluid (ACSF) consisting of (in mM): 126 NaCl, 3 KCl, 1.25 NaH_2_PO_4_, 0.6 MgSO_4_, 1.2 CaCl_2_, 24 NaHCO_3_ and 10 glucose, with neuromodulatory conditions associated with wake state modelled by inclusion of 15 µM carbachol. Extracellular recordings were obtained using micropipettes (2-5 MΩ) filled with ACSF and were bandpass filtered at 0.1 Hz to 300 Hz.

### Immunohistochemistry and confocal microscopy

For tissue preparation and immunohistochemistry mice underwent terminal anaesthesia and perfusion fixation with 4 % paraformaldehyde. Brains (or slices, following termination of electrophysiological experiments) were placed in fixative at 4 °C for >48 h and were then washed and sectioned at 20-60 µm. Control mammary tumour and naïve spleen sections were from tissue collected as part of other studies (37). The following primary antibodies were used as described previously (40): rabbit anti-Iba1 (1:500; Fujifilm Wako Chemical Corporation), mouse anti-human nuclear antigen (1:100; Merck), Alexa Fluor 647-conjugated rat anti-CD45 (1:100; Biolegend), and rabbit anti-TMEM119 (1:100; Abcam). TMEM119 staining required antigen retrieval and slices were incubated at 90 oC for 15 min in 10 mM Citrate Buffer (pH 6) containing 0.05 % Tween-20. The secondary antibodies were Alexa Fluor 488/633-conjugated goat anti-rabbit/mouse, as appropriate (1:500; Molecular Probes). Sections were counter-stained with Hoechst 33258 and mounted in VectaShield (Vector Laboratories) or Prolong Gold (Molecular Probes). Images were acquired using a Zeiss AxioObserver Z1 microscope fitted with W Plan Apochromat 20x/1.0 NA and 63x oil/1.0 NA objective lenses and LSM 880 laser scanner. Images were acquired sequentially and saved for offline processing using Zeiss Zen2 software.

### Image processing and Sholl analysis

Images were imported into the Fiji distribution of ImageJ for processing using the LSM Toolbox plugin. Brightness and contrast were adjusted using the Fiji “auto” function. Imaging depth (µm) was determined relative to that of superficial-most image for each acquisition session. Sholl analysis (41, 42) was performed on imported images as follows: First, arborisation paths on individual microglia (≤ 10 cells/image) were traced using the Simple Neurite Tracer plugin. Next, trace profiles were processed using the Sholl analysis plugin. For each cell, the number of processes intersecting concentric radii, centred on the nucleus, was recorded at 2 µm intervals and averaged to produce a Sholl plot (41). In addition, the maximum number of intersections per radius, maximum branch length (largest radius intersecting a branch), number of primary branches originating from the soma, and Schoenen ramification index (ratio of maximum number of intersections per radius and number of primary branches) were calculated (42, 43). The number of direct interactions between tumour cells and microglia (where one cell visibly touches another) was counted and divided by the number of tumour cells for a given field of view (44).

### Statistical analysis

Data are presented as mean and SEM unless stated otherwise. Statistical analysis was performed using GraphPad Prism 7. Distribution was evaluated using D’Agostino-Pearson normality test. Statistical significance was determined with Mann-Whitney tests for pairwise comparisons and ANOVA followed by Sidak’s tests or Kruskal-Wallis followed by Dunn’s tests for multiple comparisons, as appropriate. Results were considered significant at P < 0.05.

## Results

### Validation of intravital multiphoton imaging method

It was necessary to purpose-adapt our two-photon Zeiss LSM 780 microscope with an objective inverter to permit intracranial imaging. We therefore first tested the effect of the customized InverterScope on imaging performance. We confirmed that there was minimal loss of 488 nm and 920 nm excitation power exiting a Plan Apochromat 20x/0.8 objective with the InverterScope in place (Supplementary Figure 1A). There was also no impact on evenness of illumination across the field (data not shown). Using a standard fluorescent sample there was only a 15 % loss in emission at the internal detectors after 488 nm excitation and a 7 % loss in emission at the non-descanned detectors after 920 nm excitation compared to without the InverterScope (P < 0.001; Supplementary Figure 1B, C).

Given that the modified InverterScope only resulted in minimal (7 %) loss of detected emission, we next used the experimental setup to image GFP-expressing microglia in *Cx3cr1*^GFP/+^ mice. In the initial pilot study and technique development, animals were killed prior to window implantation and imaging (Figure 1A). We evaluated implantation of the optical window on the parietal bone, close to the lambdoid and sagittal sutures, as the best method for examining cellular morphology *in situ* in the outer cortex, taking into account the potential need for repeated imaging of the same animal. This approach enabled us to detect GFP-expressing microglia in *Cx3cr1*^GFP/+^ mice up to 250 µm below the window using a laser power of 6 mW (as measured at the objective) without significant loss of resolution (Figure 1Bi-iii). By increasing the laser power to 10 mW, we were able to image deeper, to depths of 350 µm, still maintaining acceptable resolution (Figure 1Biv, v). This imaging approach enabled visualisation of a highly ramified morphology with extensive arborisations, which were indistinguishable from those visualised in Iba1-stained sections using confocal microscopy (Figure 1Bvi). In order to accurately determine the effect of imaging depth on visualisation of key parameters of microglial morphology, we next quantified the extent of arborisation of microglial processes using Sholl analysis (41, 42). Analysis of microglial arborisations acquired at different imaging depths revealed typical branching profiles, equivalent to those reported previously for ramified microglia (Figure 1C) (45). There was a small shift to the left in the branching profiles for microglia imaged at depths >150 µm, indicating a slight decrease in branching with increased distance from the soma, perhaps due to a small loss in resolution and imaging sensitivity at greater depths up to 350 µm.

**Figure 1.**
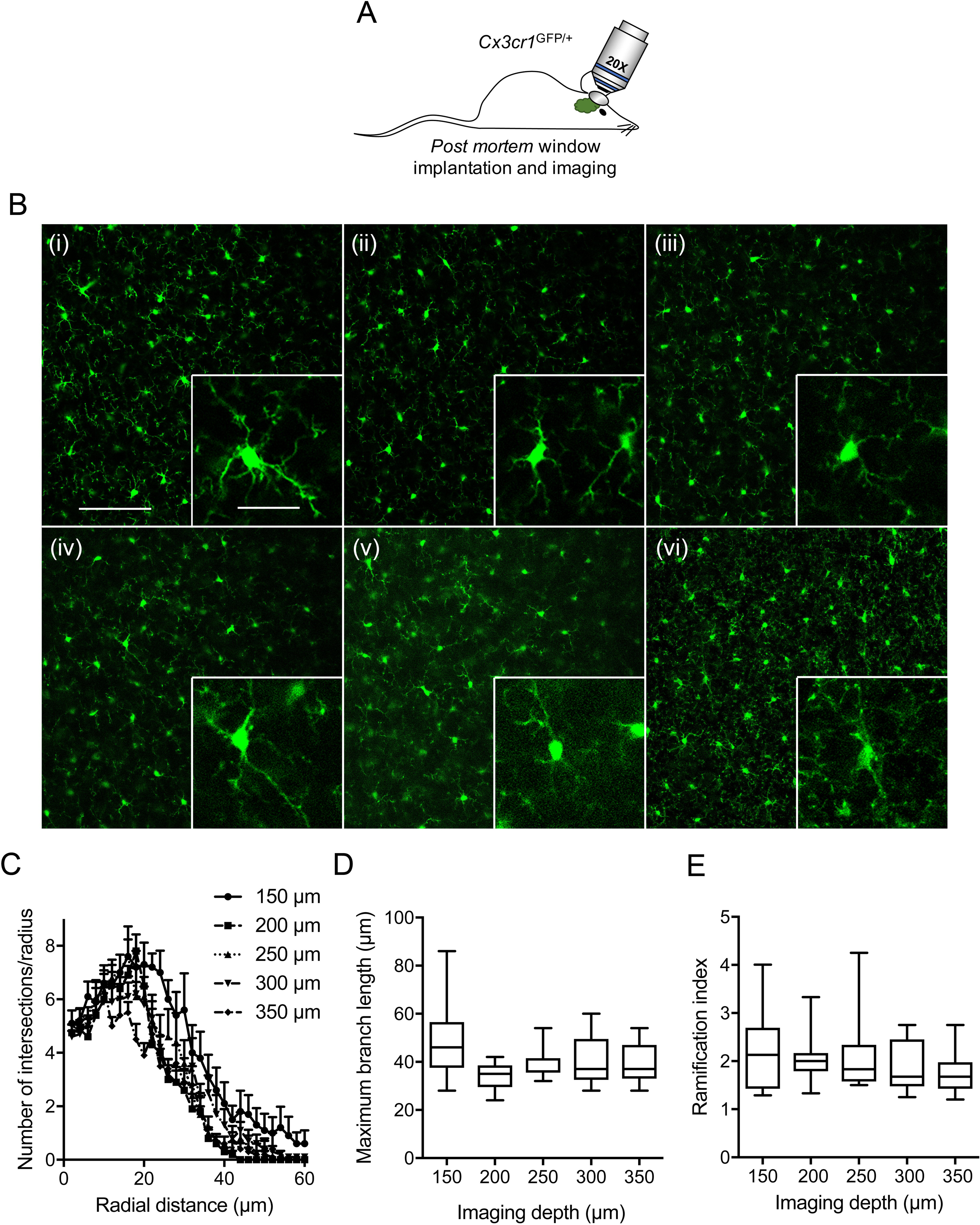
*Post mortem* visualisation of GFP-expressing microglia through a cranial imaging window. (A) *Cx3cr1*^GFP/+^ mice were euthanised, imaging window implanted as described in Methods, and microglia visualised using adapted multiphoton microscope. (B) Representative images of microglia at various distances below the imaging window: 150 µm (i), 200 µm (ii), 250 µm (iii), 300 µm (iv), 350 µm (v). (vi), microglia in a coronal section of layer 2/3 cortex from a wildtype mouse labelled with anti-Iba1 antibody. Scale bar, 100 µm. Insets: 3X magnification; scale bar, 30 µm. (C) Sholl analysis plot of microglia imaged at the indicated depths below the imaging window (n = 10 cells/group). (D) Maximum branch length (µm) of microglia imaged at the indicated depths (n = 10 cells/group). (E) Schoenen ramification index of microglia at the indicated depths (n = 10 cells/group). Laser power was 6 mW for depths 150 µm – 250 µm, and 10 mW at 300 µm and 350 µm. Box plots show median, 25^th^ and 75^th^ percentile values; whiskers are minimum and maximum values.

However, the maximum branch length (maximum radius intersecting a branch) and Schoenen ramification index were unchanged across the range of imaging depths tested (P = 0.11 and 0.43, respectively; n = 10 microglia/imaging depth; Figure 1D, E). Together, these data suggest that, using the InverterScope, it is possible to reliably detect arborisations in ramified microglia at imaging depths up to 350 µm.

### Microglia are highly ramified at steady state <u>in vivo</u>

In order to study the ramification of microglia in steady state, we next imaged GFP-expressing microglia in live anaesthetised *Cx3cr1*^GFP/+^ mice, after implantation of the optical window (Figure 2A). As with the *post mortem* imaging, we were able to visualise GFP-expressing microglia in *Cx3cr1*^GFP/+^ mice to depths up to 350 µm, maintaining acceptable resolution (Figure 2B). There was some minor loss of image quality compared to *post mortem* imaging likely as a result of pulse pressure artefacts. Second harmonic generation (SHG) imaging revealed that collagen signal was largely absent in the cortical parenchyma (data not shown), consistent with previous reports (35). Quantification of microglial arborisations using Sholl analysis revealed a typical branching pattern, similar to that calculated for *post mortem* images (Figure 2C). In addition, the maximum branch length was unchanged across the range of imaging depths tested *in vivo* (P = 0.14; n ≥ 10 microglia/imaging depth; Figure 2D). Similarly, the Schoenen ramification index was not significantly different across the same range of imaging depths (P = 0.36; n ≥ 10 microglia/imaging depth; Figure 2E). Importantly, the maximum number of intersections per radius, maximum branch length, and Schoenen ramification index were unchanged in the *in vivo* imaged animals compared to *post mortem* imaged animals (Table 1), consistent with any reduction in image quality *in vivo* not precluding detection or measurement of processes. In summary, these data suggest that visualisation of GFP-expressing microglia in anaesthetised live animals permits broadly similar measurement of microglial arborisation to *post mortem* imaging.

**Table 1.**
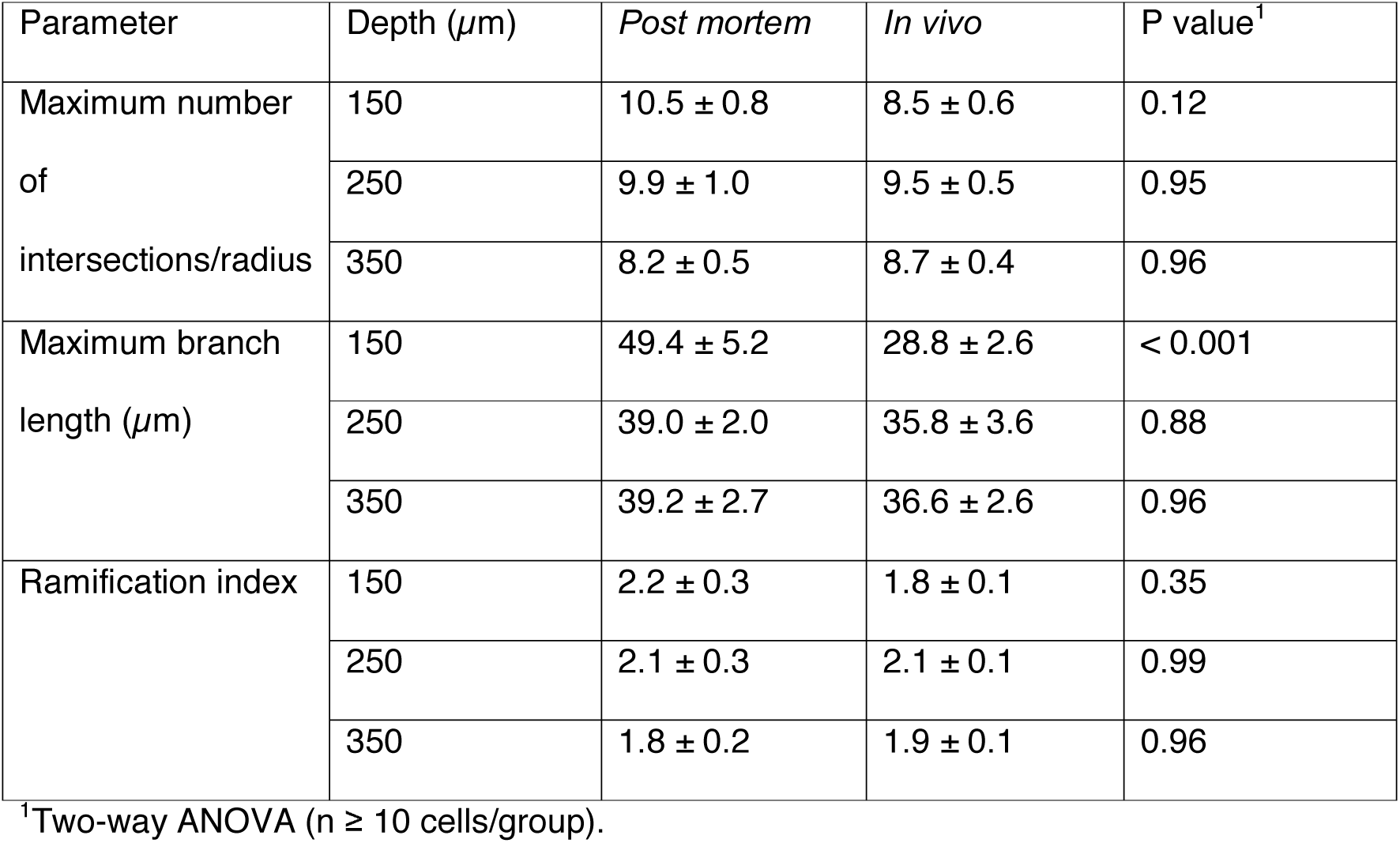
Summary of Sholl analysis results from animals imaged *in vivo* and *post mortem*.

**Figure 2.**
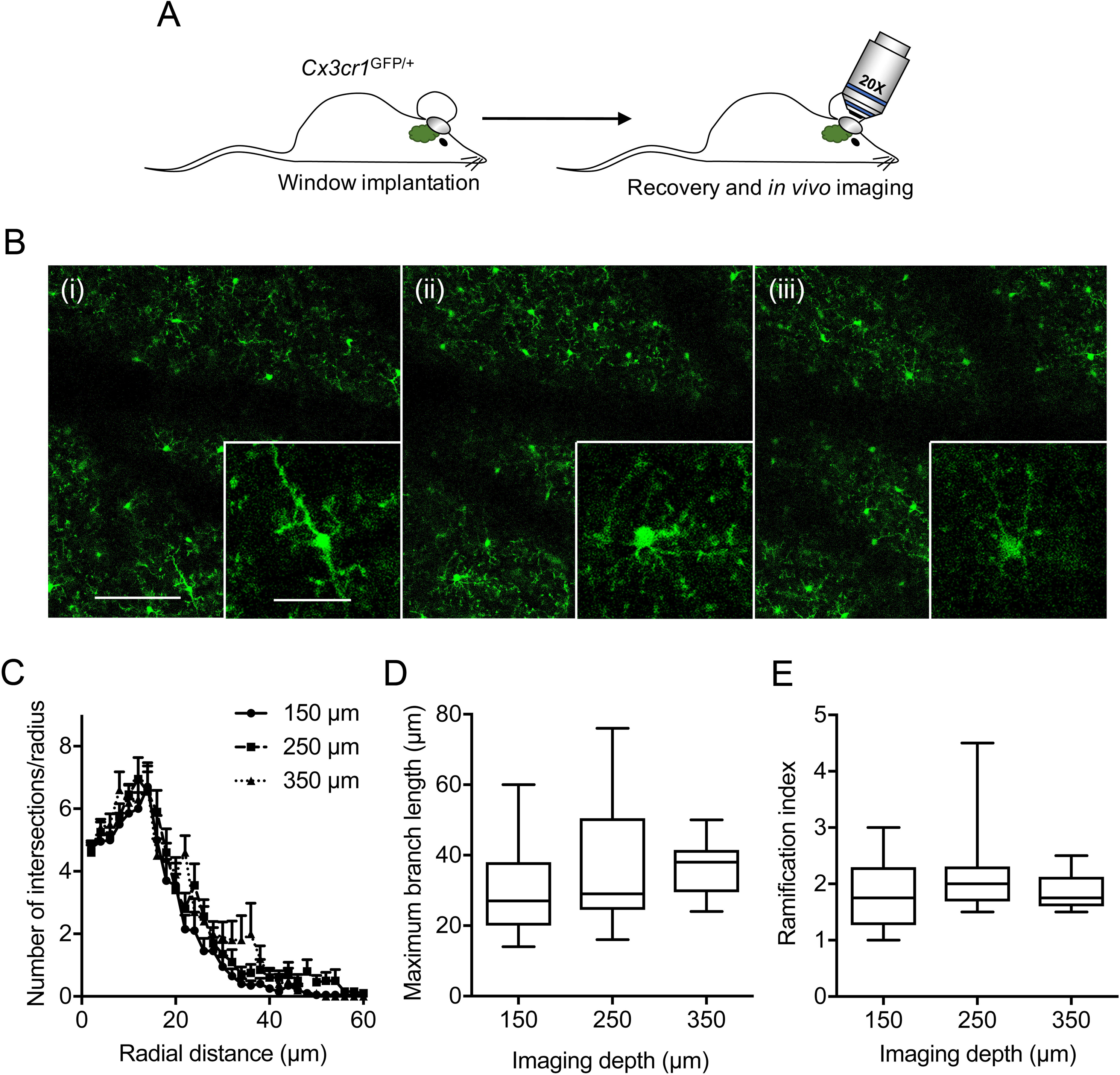
GFP-expressing microglia are ramified in healthy mice *in vivo*. (A) Imaging window was implanted in *Cx3cr1*^GFP/+^ mice as described in Methods, mice allowed to recover, and microglia visualised under anaesthesia using adapted multiphoton microscope. (B) Representative images of microglia at various distances below the imaging window: 150 µm (i), 250 µm (ii), 350 µm (iii). Scale bar, 100 µm. Insets: 3X magnification; scale bar, 30 µm. (C) Sholl analysis plot of microglia imaged at the indicated depths below the imaging window (n = 10 cells/group). (D) Maximum branch length (µm) of microglia imaged at the indicated depths (n = 10 cells/group). (E) Schoenen ramification index of microglia at the indicated depths (n = 10 cells/group). Box plots show median, 25^th^ and 75^th^ percentile values; whiskers are minimum and maximum values.

### Microglia interact with metastatic breast cancer cells

A growing body of evidence indicates that microglia promote tumour progression (20). For example, microglia have been shown to increase brain colonisation by breast cancer cells *via* a mechanism that requires Wnt signalling (46). In order to gain deeper mechanistic understanding of the processes involved, better models are needed to study microglial behaviour in the presence of metastatic cancer cells. We therefore next adapted this model to study tumour cell-microglial interactions *in vivo*. We implanted 1 × 10^4^ DsRed-expressing MDA-MB-231 metastatic breast cancer cells suspended in Matrigel on the exposed dura mater of *Cxcr31*^GFP/+^ mice, under the imaging window coverslip at time of surgery. We then imaged GFP-expressing microglia and DsRed-expressing breast cancer cells in anaesthetised mice, at various time points following recovery from surgery (Figure 3A). Using this approach, we were able to visualise invasion of tumour cells into the cortical parenchyma, together with recruitment of microglia to the lesion site at 5- and 7-days *post* implantation (Figure 3B). Consistent with this, we observed microglia surrounding and directly interacting with tumour cells (Figure 3B, arrowheads). These interactions were evident across all animals studied (additional examples are in Figure 3C). The number of microglial interactions per breast cancer cell significantly increased between 5- and 7-days post implantation (P < 0.001; n = 8 fields of view/time point; Figure 3D). In addition, the density of GFP-positive cells was significantly higher in fields of view containing tumour cells compared to animals in which no tumour cells had been implanted (P < 0.01; n ≥ 5 fields of view/condition; Figure 3E). Sholl analysis of microglial arborisations revealed that the presence of tumour cells notably reduced the number of branches compared to control animals (Figure 3F). Similarly, the maximum branch length and Schoenen ramification index were significantly reduced in the presence of cancer cells (P < 0.01 for both; n ≥ 20 microglia/condition; Figure 3G, H). Importantly, implantation of Matrigel alone without tumour cells had no effect on microglial morphology Supplementary Figure 2). Together, these data suggest that implantation of metastatic breast cancer cells induces activation and accumulation of GFP-expressing microglia.

**Figure 3.**
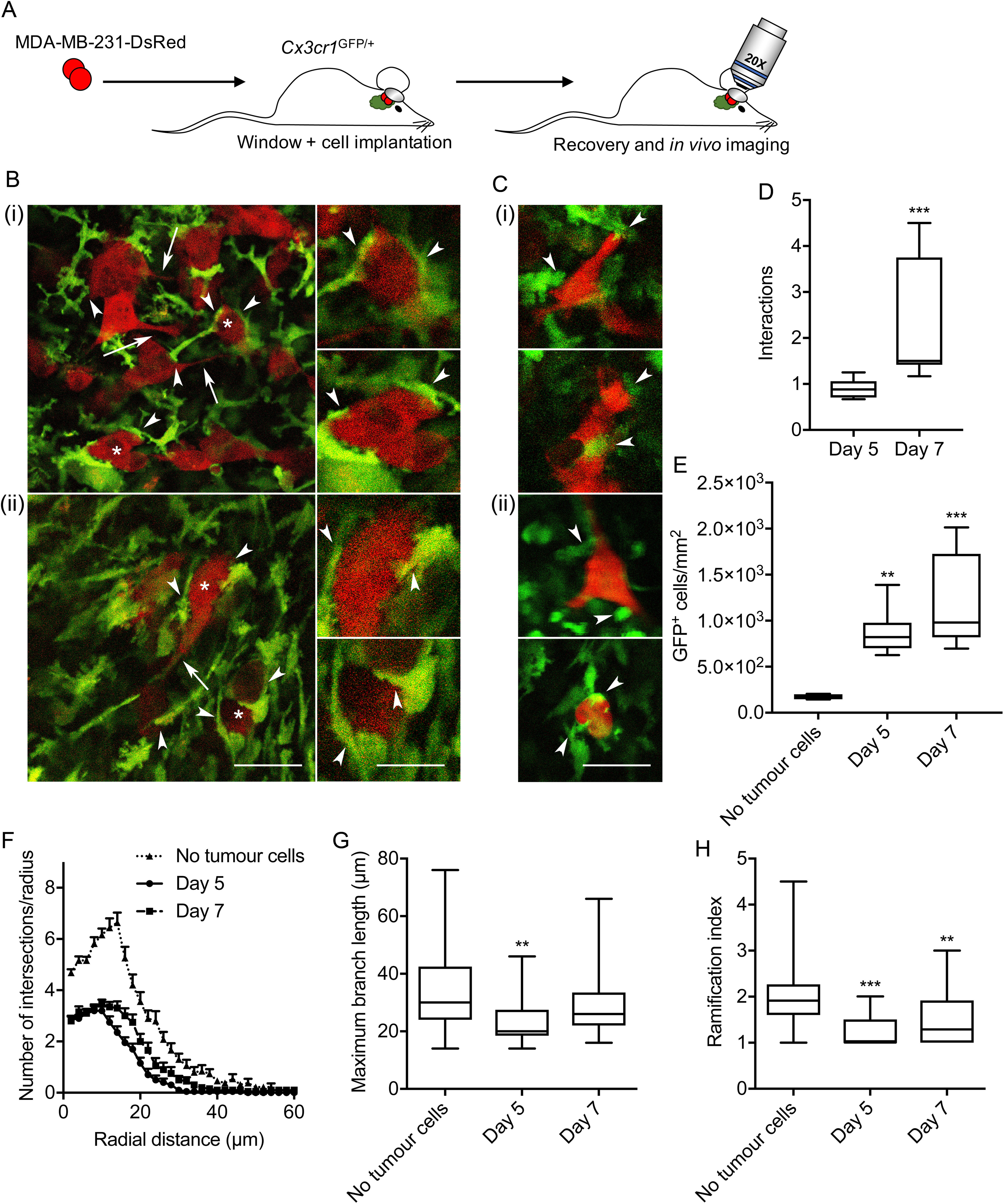
Interactions between activated microglia and metastatic breast carcinoma cells. (A) MDA-MB-231-DsRed cells were implanted under the imaging window in *Cx3cr1*^GFP/+^ mice as described in Methods, mice allowed to recover, and cells visualised under anaesthesia using adapted multiphoton microscope. (B) Representative images of activated microglia (green) and breast cancer cells (red) 5 (i) and 7 days post-implantation (ii). Asterisks in main image indicate cells in 2X magnified images on right. Scale bars, 50 µm and 25 µm, respectively. Arrowheads indicate direct interactions between microglia and tumour cells. Arrows indicate elongate neurite-like processes on tumour cells. (C) Example cellular interactions from additional mice 5 (i) and 7 days post-implantation (ii). (D) Number of microglial interactions per cancer cell 5 and 7 days post-implantation (n = 8 fields of view/time point). (E) Density of GFP-positive cells in the presence/absence of cancer cells (n ≥ 5 fields of view/time point). (F) Sholl analysis plot of microglia imaged from mice in which cancer cells had been implanted compared to mice in which no cancer cells had been implanted (n ≥ 20 cells/group). (G) Maximum branch length (µm) of microglia imaged from mice in which cancer cells had been implanted compared to mice in which no cancer cells had been implanted (n ≥ 20 cells/group). (H) Schoenen ramification index of microglia imaged from mice in which cancer cells had been implanted compared to mice in which no cancer cells had been implanted (n ≥ 20 cells/group). Box plots show median, 25^th^ and 75^th^ percentile values; whiskers are minimum and maximum values. **P < 0.01; ***P < 0.001; Mann-Whitney test for (D) and Kruskal-Wallis with Dunn’s post hoc test for (E), (G) and (H).

### Persistent alteration of tissue architecture and electrical activity following tumour cell implantation

The accumulation of microglia to the lesion site, which started from 5 days, was maintained at 9- and 13-days post implantation (Figure 4A-C). In particular, there was notable accumulation of GFP-positive cells at the periphery of the lesion (Figure 4A, arrows). Interestingly, however, we were no longer able to detect DsRed-positive tumour cells at 9 and 13 days, suggesting that the microglial-induced immune response was capable of clearing the tumour cells from the lesion site (Figure 4B, C). In agreement with this observation, *ex vivo* confocal imaging of tissue sections at the end of the experiment revealed that although some DsRed-expressing tumour cells were still present superficially at the pia mater, none were detectable deeper into the tissue (Figure 5A, B, arrowheads). Similarly, sections stained with an anti-human nuclear antigen antibody showed an absence of tumour cells at the lesion site (Figure 6). However, we were able to detect punctate acellular DsRed signal adjacent to, or within some GFP-expressing cells, consistent with the possibility that the microglia had destroyed and engulfed some of the tumour cells (Figure 5A-C, arrows). In addition to microglia, CX3CR1 is also expressed on macrophages, certain subtypes of T cells and natural killer cells (38, 47, 48), raising the possibility that some of the GFP-positive population at the lesion site may constitute other cell types. The microglia-specific marker TMEM119 (49) was expressed on ramified microglia in control brain parenchyma and there was also strong staining at the lesion site (Figure 7A, B). There was some limited infiltration of CD45^hi^ GFP^−^ leukocytes at the lesion site compared to normal brain parenchyma (Figure 8A, B). However, the majority of GFP^+^ cells were CD45^low or −^, consistent with these cells being microglia (Figure 8B). Together, these data suggest that the GFP^+^ cells at the lesion are predominantly microglia. In addition, the lack of CD45^hi^ cells suggests that there is minimal infiltration of other immune cells in response to the xenograft.

**Figure 4.**
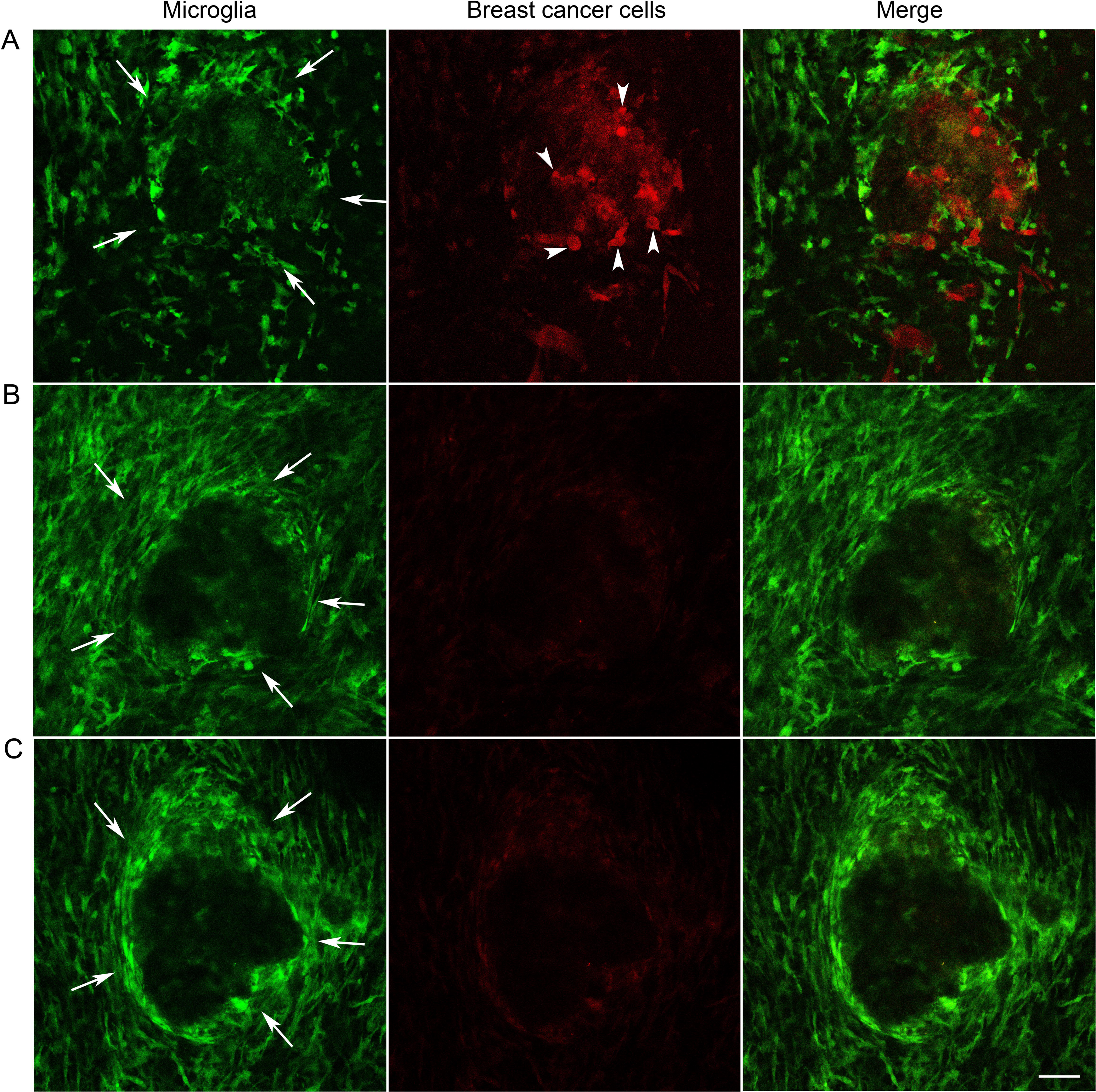
Persistent accumulation of microglia at the tumour site for 13 days following implantation. Microglia (green) and tumour cells (red) were visualised under anaesthesia using adapted multiphoton microscope at 5 (A), 9 (B), and 13 (C) days following implantation. Scale bar, 50 µm. Arrows indicate periphery of lesion where microglia have accumulated. Arrowheads indicate tumour cells within lesion, which are no longer visible after 9 days.

**Figure 5.**
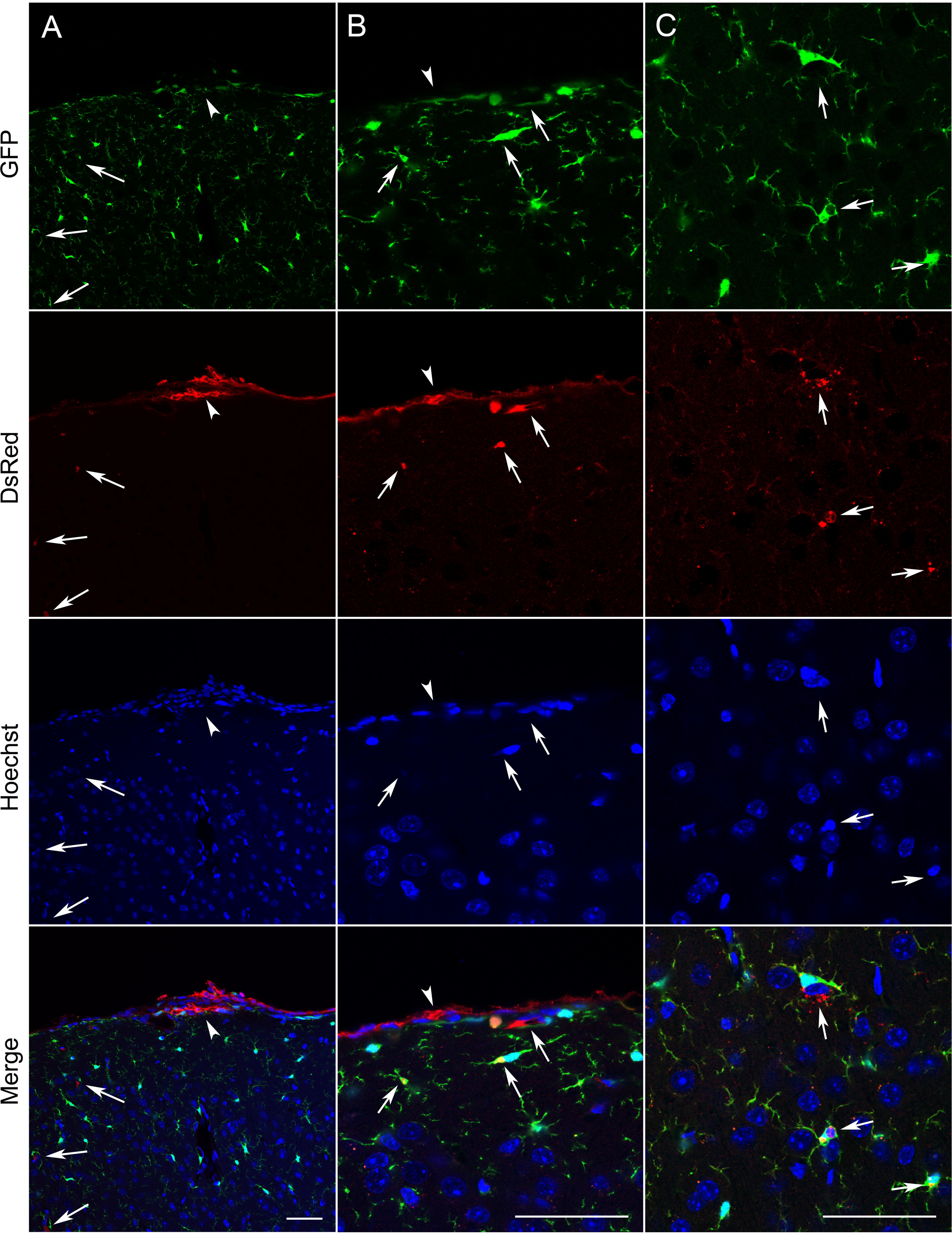
Tumour cells are retained superficially but not deeper into the brain parenchyma. Confocal images at 20x (A) and 63x (B, C) of microglia (green), tumour cells (red) and nuclei (blue) in coronal sections obtained from *Cx3cr1*^GFP/+^ mice killed 22 days following implantation of MDA-MB-231-DsRed cells under the imaging window. Scale bars, 50 µm. Arrowheads indicate presence of viable DsRed-expressing tumour cells on outer surface of pia mater. Arrows indicate DsRed signal adjacent to, or within, GFP-expressing microglia deeper into brain parenchyma.

**Figure 6.**
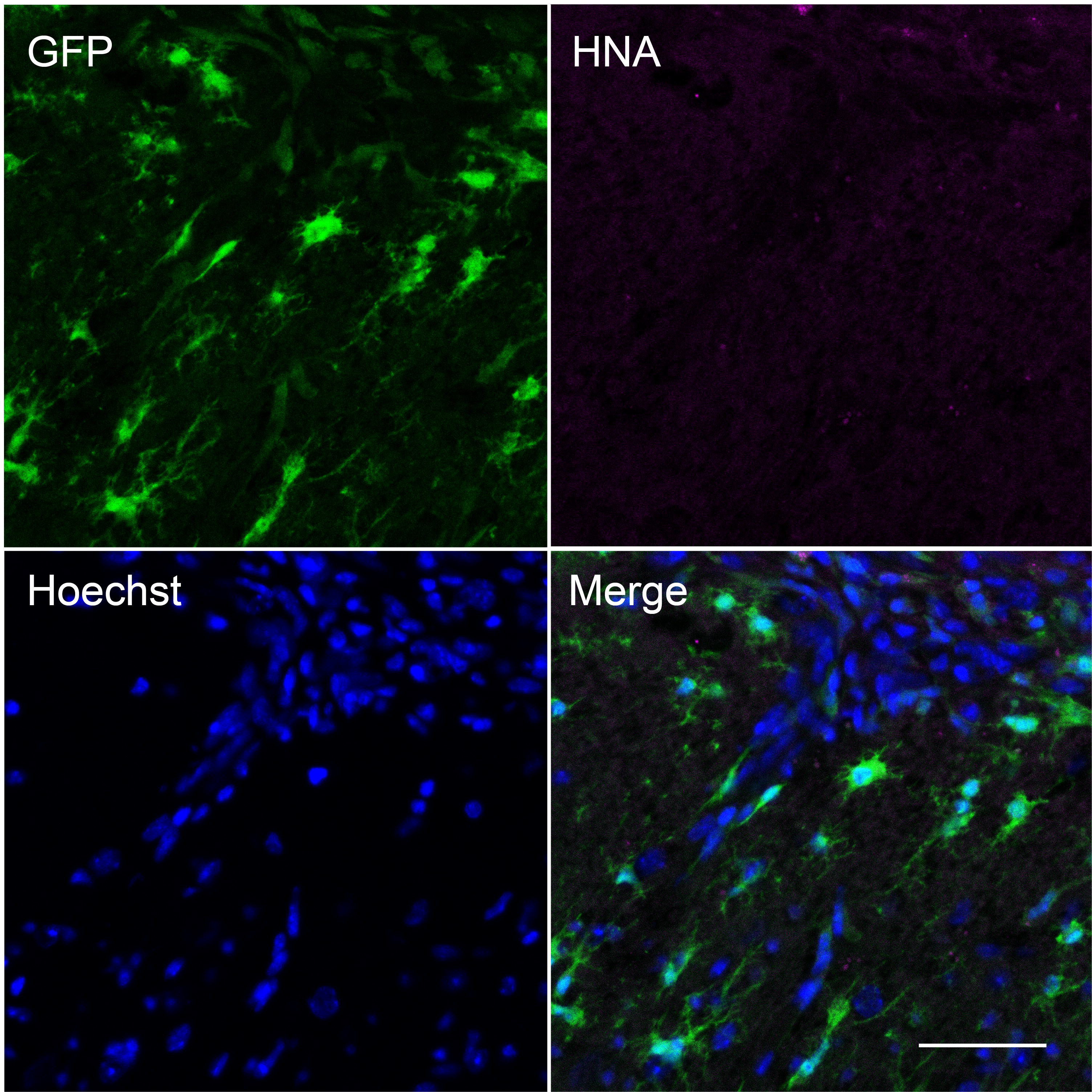
Absence of human nuclear antigen-positive cells at the lesion site. Confocal image at 20x of microglia (green), human nuclear antigen (HNA; magenta) and nuclei (blue) in coronal section obtained from a *Cx3cr1*^GFP/+^ mouse killed 6 days following implantation of tumour cells under the imaging window. A positive control MDA-MB-231 orthotopic mammary xenograft tumour section showed strong HNA staining (data not shown). Scale bar, 50 µm.

**Figure 7.**
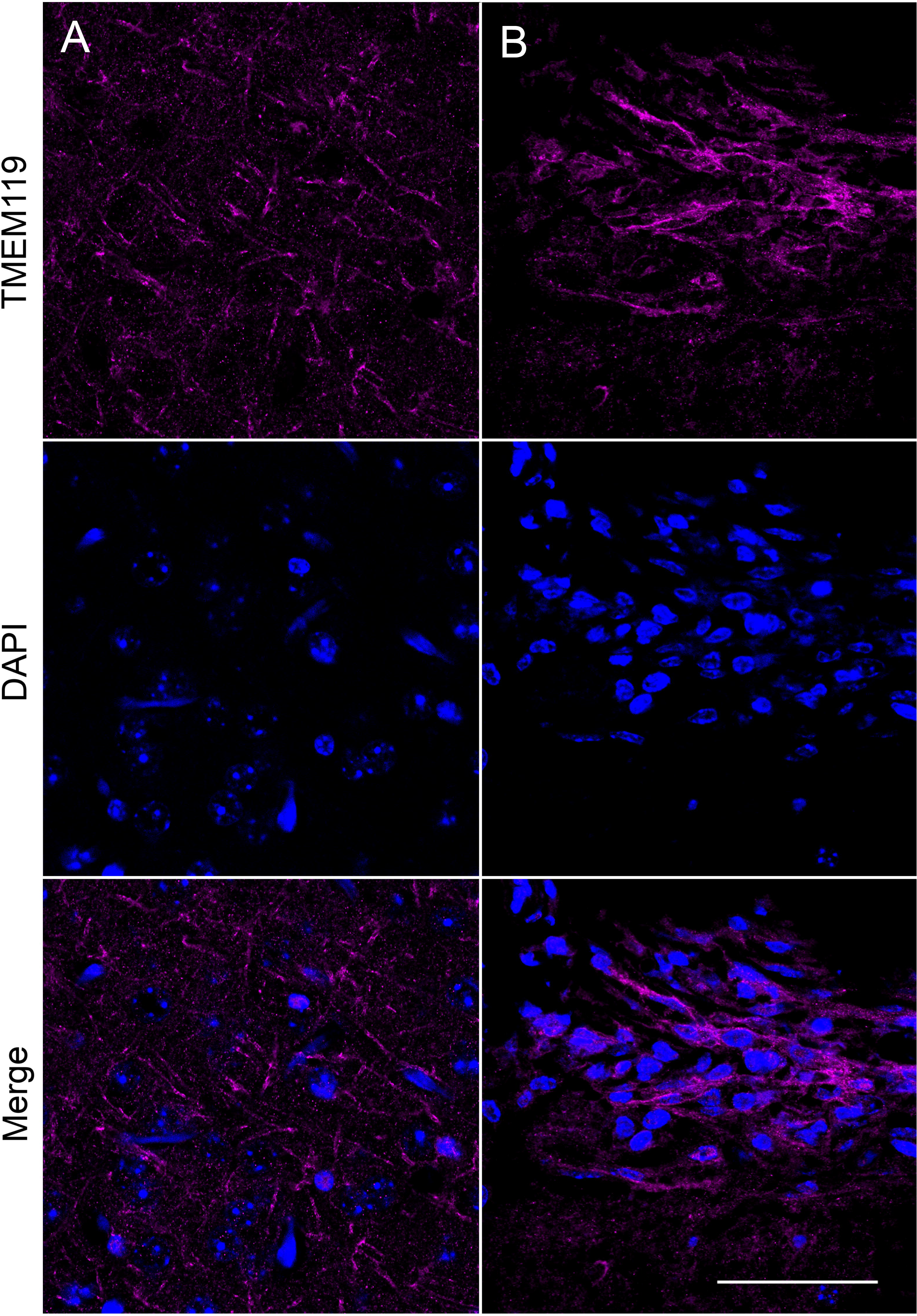
Presence of TMEM119-positive microglia at the lesion site. Confocal images at 63x of TMEM119-positive microglia (magenta) and nuclei (blue) in coronal section obtained from a *Cx3cr1*^GFP/+^ mouse killed 6 days following implantation of tumour cells under the imaging window. (A) Control parenchyma away from lesion site. (B) Lesion site underneath location of imaging window. Scale bar, 50 µm.

**Figure 8.**
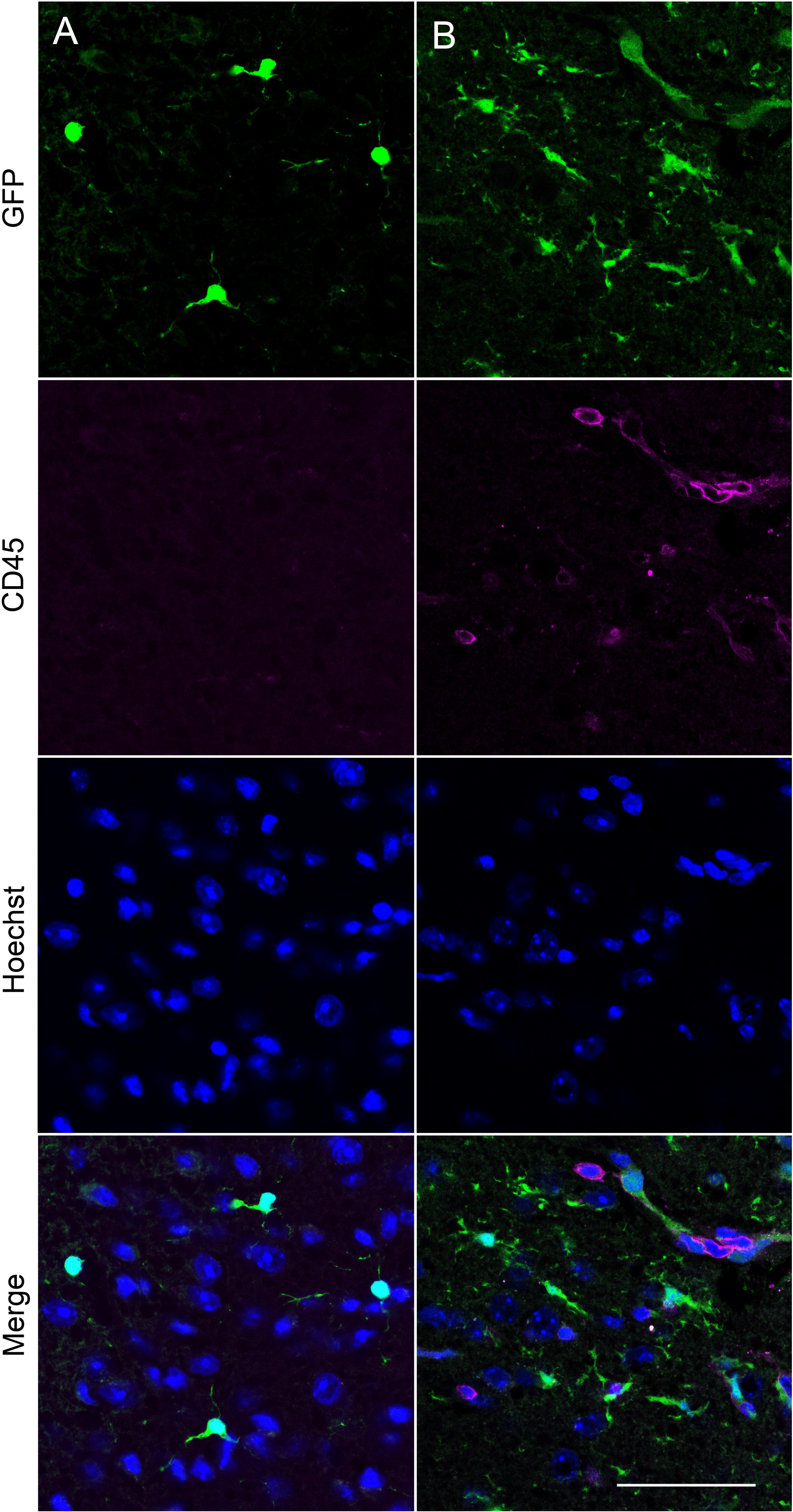
Infiltration of CD45^hi^ cells at the lesion site. (A, B) Confocal images at 63x of GFP-positive microglia (green), CD45 (magenta) and nuclei (blue) in coronal section obtained from a *Cx3cr1*^GFP/+^ mouse killed 6 days following implantation of tumour cells under the imaging window. (A) Control parenchyma away from lesion site. (B) Lesion site underneath location of imaging window. A positive control naïve spleen section showed strong CD45 staining (data not shown). Scale bar, 50 µm.

Given the striking observation of significant and persistent microglial accumulation to the tumour site, we postulated that the lesion may disrupt normal cortical electrophysiological function. We therefore next evaluated spontaneous electrical activity in coronal tissue slices under the imaging window and on the contralateral side to the implantation site. Spontaneously-occurring local field potential spike events were observed in slices taken from the region of the brains underlying the tumour cell implantation sites in all cases. Events were only seen in superficial layers, particularly layer 2 (L2) and never seen in deep layers (Figure 9A) or in any layer contralateral to the inoculation site. Mean spike amplitude was 98 ± 6.0 µV (n = 138 events from slices from 2 mice) with 5/138 outliers where amplitude was >0.5 mV. Field spike durations were highly variable and non-normally distributed (Figure 9B). Three modal peak halfwidths were seen at 48, 111 and 215 ms. Morphology of the field spikes was highly dependent on recording electrode position relative to the lesion (Figure 9C). Within the lesion itself bursts of multi-unit activity occurred with no concurrent field spike. This pattern was reversed peri-annularly around the lesion wall in L2, with overt field spikes seen with no concurrent multi-unit activity. In summary, the imaging and electrophysiological data reveal that in this model there is a significant and persistent microglial response to the location of metastatic breast cancer cells, and although this is capable of removing some of the tumour cells, there remains significant disruption to normal electrical activity within and adjacent to the lesion site.

**Figure 9.**
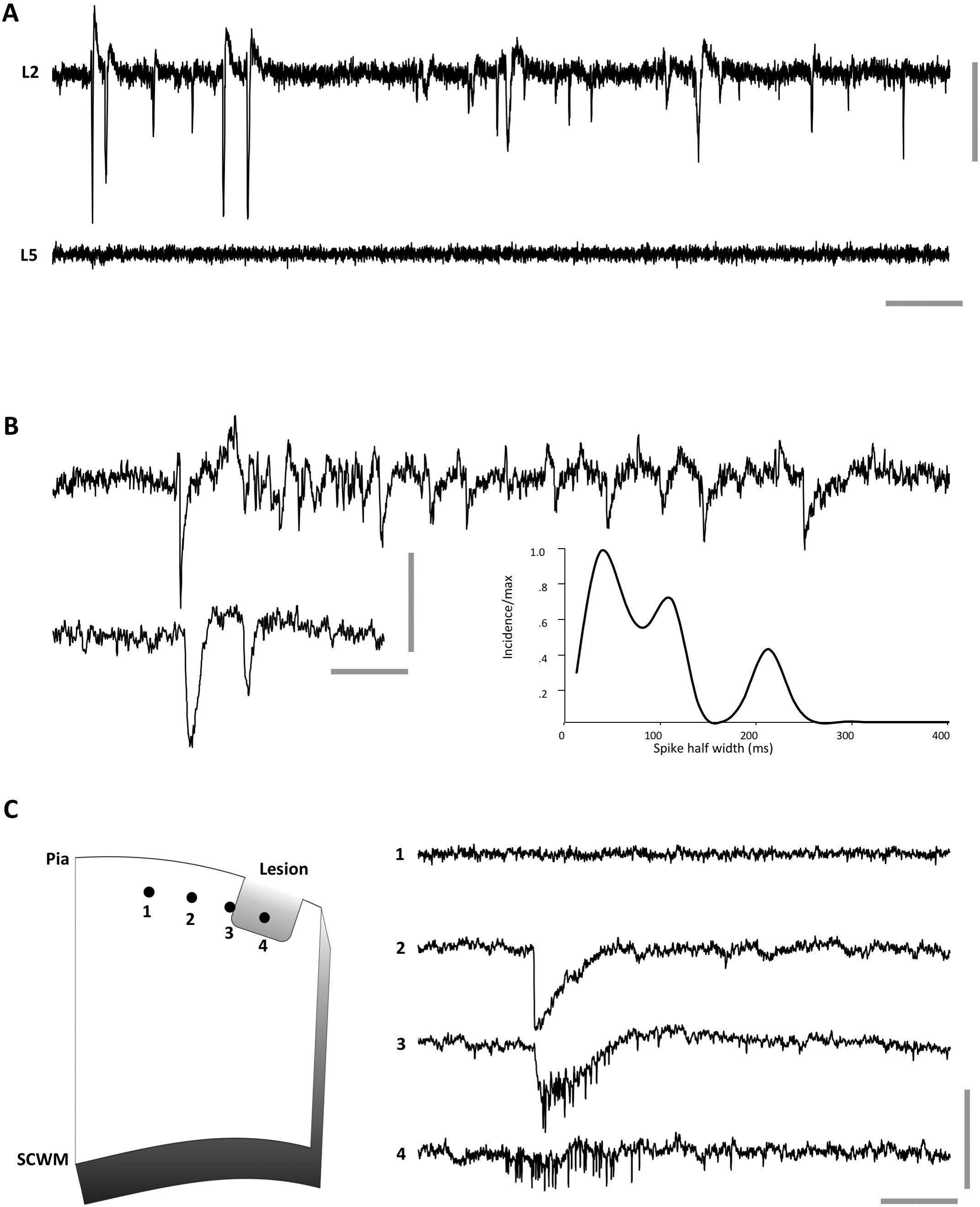
Basic properties of lesion-associated extracellular spikes. (A) 25 s example of spontaneous, transient spike generation in superficial layers (cortical layer 2, L2) on the side of the brain ipsilateral to the tumour cell implantation site. Note concurrent recordings from deep (cortical layer 5, L5) layers demonstrated no spiking. (B) Example L2 recordings demonstrating the variability of spike generation. Illustrated are an example of the most common discharge – pairs of field spikes or relatively broad halfwidth and an example of a narrow halfwidth initial spike leading to a polyspike-like pattern of afterdischarge. Graph shows distribution of initial spike halfwidths (normalised to maximum incidence) for 138 spontaneous events from n=4 slices from N=2 mice. Note the multimodal nature of the distribution of spike widths. (C) Spikes were highly localised annularly around the radial walls of the lesions in L2. Cartoon illustrates four recording sites at different positions from the core of the lesion in L2. Example traces taken from each of these positions reveal multi-unit activity only within the body of the lesion, multiunit and field spike at the border of the lesion and field spike only extending <200 µm tangentially from this. Scale bars 0.1 mV, 2 s (A), 0.5 s (B), 0.1 s (C).

## Discussion

In this study, in mice expressing GFP under control of the endogenous *Cx3cr1* promoter, we were able to visualise microglial behaviour in the cortex at a depth of up to 350 µm using intravital imaging without significant loss of resolution. Sholl analysis revealed that the arborisation of resting microglia was unchanged across the range of imaging depths tested, both *ex vivo* and *in vivo*. Implantation of DsRed-expressing MDA-MB-231 breast cancer cells under the optical window resulted in recruitment and activation of microglia to the site of metastatic cancer cells. The immune response cleared the majority of the tumour cells within 9 days, but the accumulation of microglia to the lesion persisted for at least 13 days. This significant cortical disorganisation gave rise to aberrant spontaneously-occurring local field potential spike events within the tumour and peri-annularly in superficial layers.

Intravital imaging of microglial morphology enables the study of cellular changes in the native microenvironment in real-time and thus is a critical approach to study behavioural changes in response to injury or infection (14, 36). Although it is possible to culture microglia *in vitro*, when removed from the brain, their behaviour changes (50) and so intravital imaging provides a useful means to study microglial arborisation in living cells *in situ*. The microglial behaviour captured here using intravital imaging broadly agrees with previous reports (14, 36), together with morphological data recorded from brain tissue sections using immunohistochemistry (45). Quantification of microglial morphological characteristics using Sholl analysis reveals broadly similar arborisation profiles and Schoenen ramification indices to those reported in tissue sections (41, 42, 45), underscoring that there was no activation of microglia in the parenchyma under the imaging window as a result of the surgery or implantation of Matrigel alone. Furthermore, the high degree of consistency between our intravital imaging data and the reported *ex vivo* immunohistochemistry data (45) suggest that any loss of resolution as a result of imaging *in vivo* was minimal.

Our data demonstrate a remarkable change in microglial morphology in response to the presence of metastatic breast cancer cells. We observed accumulation of microglia to the lesion site, together with significant reduction in ramification, consistent with acquisition of an activated phenotype. To our knowledge, this is the first time that intravital brain imaging has been used to visualise the behaviour of microglia in the presence of metastatic breast cancer cells. The implantation site represents a leptomeningeal metastasis, which accounts for approximately 11-20 % of brain metastases (5). However, we also found significant infiltration of tumour cells into the outer cortical parenchyma and recruitment of microglia to the lesion, suggesting that the model may have broader utility for understanding tumour cell-microglial interactions in parenchymal brain metastases.

As CX3CR1 is also expressed on peripheral blood monocytes, natural killer cells and dendritic cells in addition to microglia (38, 47), it is possible that some of the accumulation of GFP-positive cells that we observed at the lesion site may be due to recruitment of other CX3CR1-expressing immune cells. However, the TMEM119 staining argues against this and suggests that there was minimal infiltration of other immune cell types. The data from our model agree with previous observations from mice and patients that activated microglia cluster around brain metastases (20), suggesting that the majority of the peritumoral GFP-positive cells are indeed microglia. We found that the accumulation of microglia persisted for the 13-day duration of the experiment following implantation of the breast cancer cells. However, the number of intact breast cancer cells was dramatically reduced after 5 days, although we could still detect isolated DsRed-positive cells and punctate acellular DsRed fluorescence at the lesion site at the end of the experiment. This rapid clearance of the tumour cells may reflect, in part, a graft versus host immune response to the xenotransplantation of a human breast cancer cell line into an immunocompetent mouse (51), although there was very limited infiltration of CD45^hi^ immune cells at the lesion site. Nonetheless, the results presented here suggest that imaging of microglia in the context of metastatic breast cancer cells in the brain may yield novel insights into their functional contribution to tumour progression (20). In particular, intravital imaging approaches such as this have the advantage of enabling visualisation of cellular changes in real-time using a system that is easy to manipulate (14, 30, 36).

Primary and metastatic brain tumours can have a range of neurological consequences such as headaches, mild cognitive impairment, and epileptic seizures (12). This aspect of the disease burden in the brain has a significant impact on quality of life and is difficult to treat as the mechanistic basis is poorly understood (10-12). Several mechanisms have been proposed to explain the presentation of tumour-associated epilepsy, including disrupted glutamate homeostasis, altered ion channel expression, altered network activity, and inflammation (52-55). We found aberrant spontaneously-occurring local field potential spike events local to the tumour. In addition, the activity pattern differed within the lesion compared to peri-annularly around the lesion wall, suggesting that changes within the tumour and peritumoral tissue may differentially mediate altered electrical activity. The altered excitability and electrical signatures observed here resemble interictal-like epileptic wave forms that have been associated with poor cognitive performance and have been seen in electroencephalograms (EEGs) of patients presenting with epilepsy (56), suggesting that our model may recapitulate early stages of cortical disorganisation that could subsequently lead to tumour-associated epilepsy. Furthermore, the data raise the possibility that peritumoral microglial activation and accumulation may play a critical role in local tissue changes leading to aberrant cortical activity.

## Conclusions

The results presented here show that it is possible to visualise the microglial response to metastatic breast tumour cells using intravital imaging. Using this approach, we demonstrate that activated microglia accumulate at the lesion site, giving rise to peritumoral cortical disorganisation. Although the immune response rapidly cleared the majority of tumour cells in this model, there was aberrant spontaneous activity both within the lesion and peritumorally. Further work is required to establish the role of microglial activation in the generation of altered excitability and seizure activity in the context of breast cancer metastasis to the brain.

## Declarations

### Ethics approval and consent to participate

Animal procedures were performed following approval from the University of York Animal Welfare and Ethical Review Body and under authority of a UK Home Office Project Licence. Surgical procedures were performed within the regulations of the UK Animals (Scientific Procedures) Act, 1986.

### Consent for publication

Not applicable.

### Availability of data and materials

The datasets used and/or analysed during the current study are available from the corresponding author on reasonable request.

### Authors’ contributions

AS, MY, PK, MW, SC and WB contributed to the conception and design of the work. AS, MY, JM, AJ, PO’T PK, MW, SC and WB contributed to acquisition, analysis, and interpretation of data for the work. AS, MY, PK, MW, SC and WB contributed to drafting the work and revising it critically for important intellectual content. All authors approved the final version of the manuscript.

## Acknowledgements

We thank Prof V. DePaola (Imperial College, UK) for help in establishing the intravital imaging protocol.

