## Supplementary figures and images for "Metastatic breast cancer cells induce altered microglial morphology and electrical excitability *in vivo*"

### Supplementary Figure 1

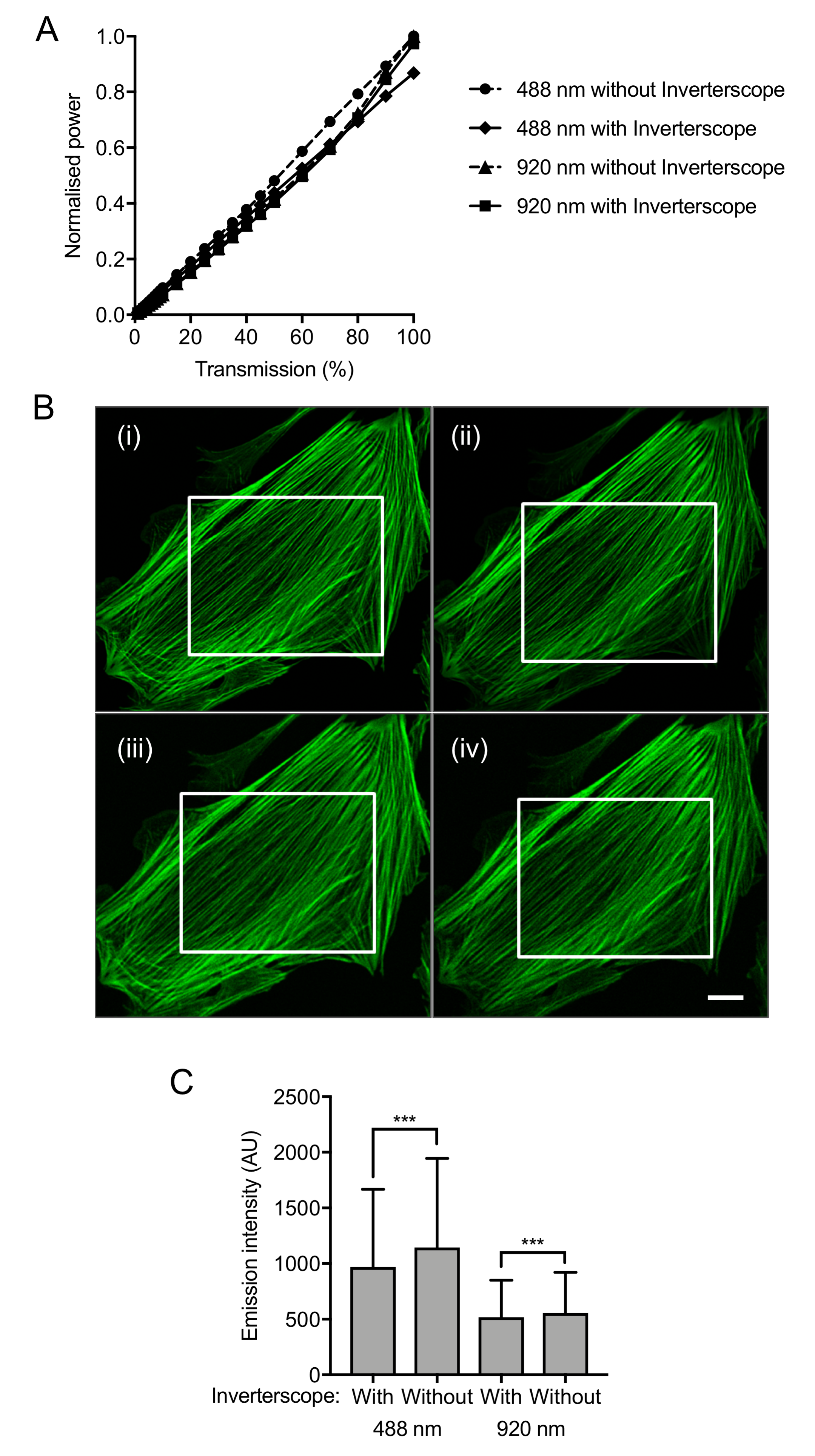

### Supplementary Figure 2

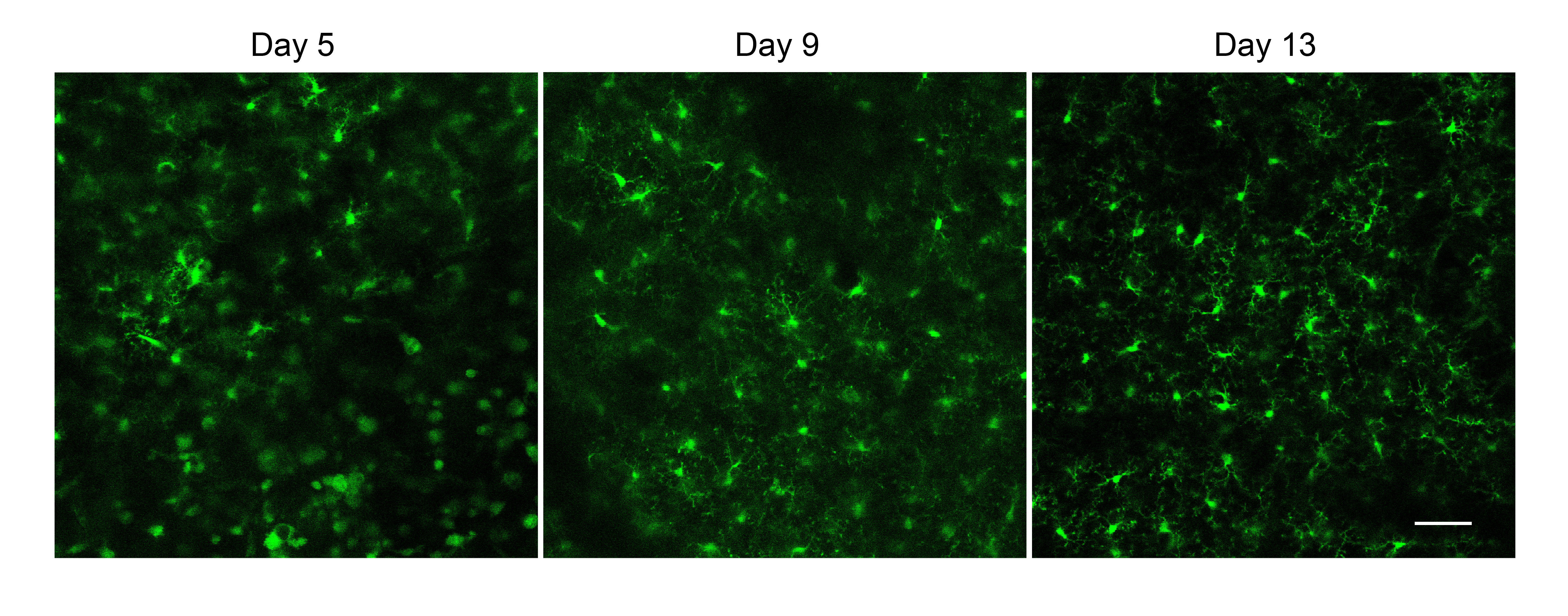
