## Supplemental Data 1 for "Metastatic breast cancer cells induce altered microglial morphology and electrical excitability *in vivo*"

**Supplementary Figure 1.** Effect of InverterScope on laser power and emission intensity. (A) Laser power at 488 nm and 920 nm exiting the objective with and without the InverterScope, measured at the indicated transmission percentages using a Coherent Fieldmate power meter. (B) AlexaFluor 488 phalloidin emission intensity in a standard sample imaged with and without the InverterScope: (i, ii) 488 nm excitation (40 mW) with 500-550 nm emission directed to the internal detectors. (iii, iv) 920 nm 2-photon excitation (13 mW) with 500-550 nm emission directed to the non-descanned detectors. i,iii show a cell without the InverterScope and ii,iv show the same cell with the InverterScope. (C) Mean intensity measured within a defined region of interest (white box). ***P < 0.001, ANOVA with Sidak’s multiple comparisons test. Scale bar, 20 µm. Data are mean ± SD.

**Supplementary Figure 2.** Lack of effect of Matrigel on microglial morphology. Microglia (green) were visualised under anaesthesia using adapted multiphoton microscope at 5, 9, and 13 days following implantation of Matrigel alone without tumour cells. Scale bar, 50 µm.
